# Visual Entrainment at 10 Hz causes periodic modulation of the Flash Lag Illusion

**DOI:** 10.1101/515114

**Authors:** Samson Chota, Rufin VanRullen

## Abstract

It has long been debated whether visual processing is, at least partially, a discrete process. Although vision appears to be a continuous stream of sensory information, sophisticated experiments reveal periodic modulations of perception and behavior. Previous work has demonstrated that the phase of endogenous neural oscillations in the 10 Hz range predicts the “lag” of the flash lag effect, a temporal visual illusion in which a static object is perceived to be lagging in time behind a moving object. Consequently, it has been proposed that the flash lag illusion could be a manifestation of a periodic, discrete sampling mechanism in the visual system. In this experiment we set out to causally test this hypothesis by entraining the visual system to a periodic 10 Hz stimulus and probing the flash lag effect (FLE) at different time points during entrainment. We hypothesized that the perceived FLE would be modulated over time, at the same frequency as the entrainer (10 Hz). A frequency analysis of the average FLE time-course indeed reveals a significant peak at 10 Hz as well as a strong phase consistency between subjects (N=26). Our findings provide evidence for a causal relationship between alpha oscillations and fluctuations in temporal perception.

## 1. Introduction

It has been suggested that perception may be a periodic process (VanRullen, 2016). The detection probability of near-threshold stimuli has been shown to oscillate at around 10 Hz in vision (Busch, Dubois, & VanRullen, 2009). Reaction times are governed by similar periodic fluctuations in the 10 Hz range (Callaway & Yeager, 1960; Başar, Başar-Eroglu, Karakaş, & Schürmann, 2001). While it was theorized that the endogenous alpha oscillations of the brain might give rise to these periodicities, their functional relevance is an ongoing enigma. One possible functional role for these neural rhythms is that of a periodic sampling mechanism (Busch et al., 2009; Haegens, Nácher, Luna, Romo, & Jensen, 2011; Lőrincz, Kékesi, Juhász, Crunelli, & Hughes, 2009; Samaha & Postle, 2015; VanRullen, 2016; VanRullen & Koch, 2003; Vijayan & Kopell, 2012): in order to reduce the complexity of the visual stream, the brain is repeatedly dividing the incoming visual information into temporal chunks or windows; these chunks are then passed on for further processing and subsequently made consciously available. Evidence for rhythmic fluctuations in perception is abundant (VanRullen, 2016). It was repeatedly shown that the phase of 5-15Hz oscillations has an influence on the efficiency of stimulus processing, mostly by providing correlational evidence via EEG (Ai & Ro, 2013; Busch et al., 2009; Dugué & VanRullen, 2017; Mathewson, Gratton, Fabiani, Beck, & Ro, 2009). These findings are supported by causal evidence from neural entrainment studies. Periodic stimulation at 10 Hz interacts with the endogenous alpha rhythm, which is a prime candidate for the brain mechanism for visual sampling. Crucially, the entrained oscillations were behaviorally significant, influencing the rate of target detection and the visibility of masked stimuli (Keitel, Quigley, & Ruhnau, 2014; Mathewson, Fabiani, Gratton, Beck, & Lleras, 2010; Mathewson et al., 2011; Romei, Gross, & Thut, 2010; Spaak, Lange, & Jensen, 2014; Thut et al., 2011). However, while these findings show clear periodicities in perception, they might be simply explained by fluctuations in neural excitability, and thus may not provide direct evidence for a true discreteness of visual perception. In order to prove the latter, one needs to show a rhythmic modulation of the *temporal parsing* of events, i.e. a periodicity in time perception itself. These two notions, rhythmic fluctuations of excitability versus rhythmic fluctuations of temporal perception, can be thought of as “soft” and “hard” versions of the discrete perception idea (VanRullen, 2016). The “soft” version claims that it is merely the excitability state of the brain that is oscillating, leading to fluctuations in detection probabilities or perceived intensities. The “hard” version goes one step further and claims that within these windows, time perception is impossible. In other words, according to the “hard” version the visual system can be imagined as a biological camera with a certain framerate, averaging collected information within one frame. Few studies have explicitly tested the periodicity of *temporal* perception (and thus, the more conservative or “hard” definition of discrete perception). Samaha and Postle demonstrated that the individual alpha peak frequency is predictive of the performance in the two-flash-fusion paradigm (Samaha & Postle, 2015). Opposite phases of the alpha rhythm are related to perception of synchronicity and a-synchronicity respectively (Milton & Pleydell-Pearce, 2016; Valera, Toro, Roy John, & Schwartz, 1981). Of high importance for this study are the findings by Chakravarthi and VanRullen (2012), relating the perceived flash lag duration to the phase of endogenous alpha oscillations using EEG.

In this study we were interested in providing *causal* evidence for a periodic modulation of time perception. To our knowledge only one study has succeeded in this so far. In a recent publication by Ronconi et al. (Ronconi, Busch, & Melcher, 2018) it was shown that visuo-auditory entrainment at the individuals alpha frequency + or – 2 Hz was able to modulate the integration or segregation of two stimuli in close temporal proximity. In this study we seek to provide an important addition to these findings by demonstrating a causal influence of alpha on the perception of time, using a well investigated visual illusion, the flash lag effect (FLE).

Discrete sampling in the visual system has previously been hypothesized to underlie the FLE (Chakravarthi & Vanrullen, 2012; Schneider, 2018). In the FLE a stationary object is shortly presented (“flashed”) alongside a moving object. Although the position of both objects is identical at the onset of the flashed, stationary object, observers systematically judge the flashed object to be lagging behind (Figure 1A).

**Figure 1.**
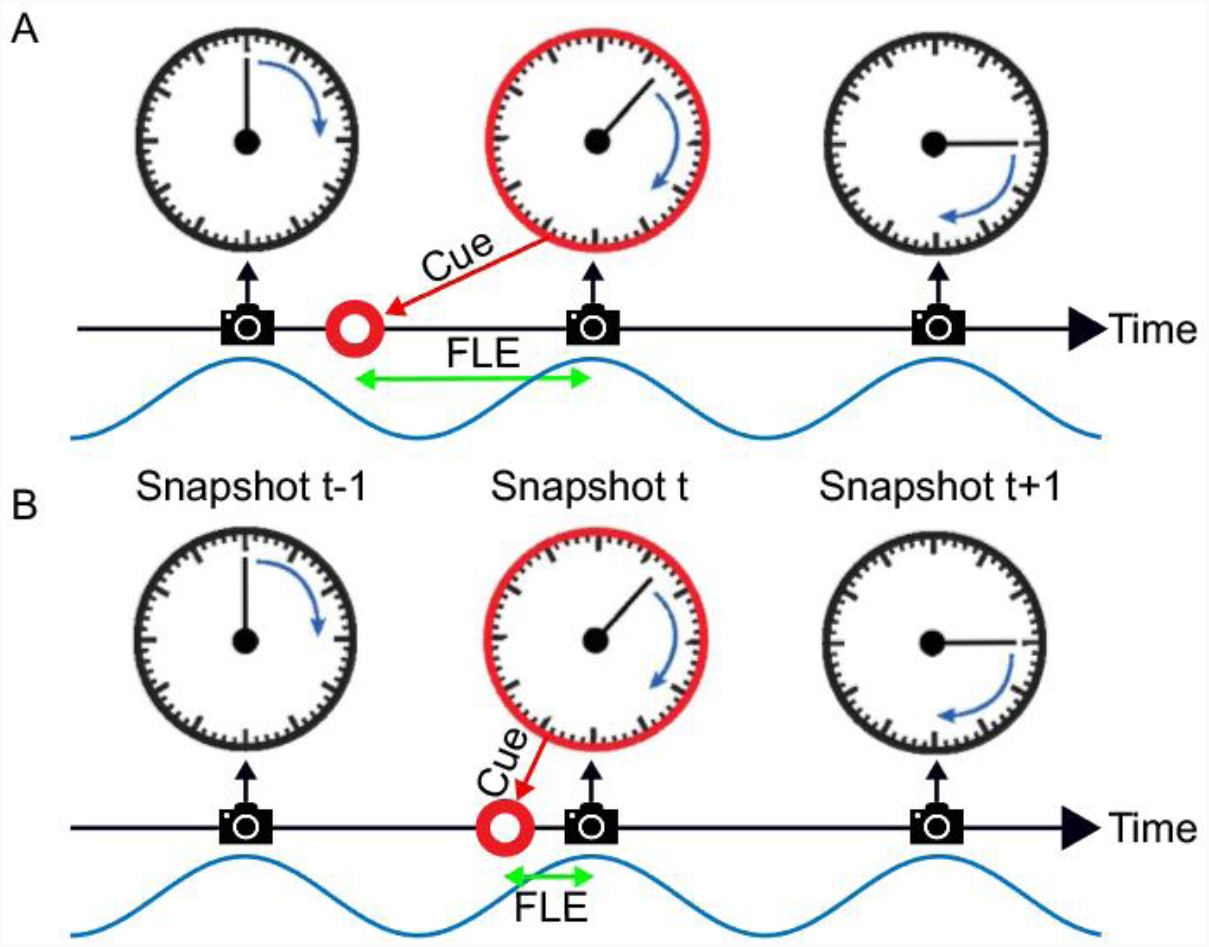
The flash lag illusion as a consequence of discrete sampling. Following the hypothesis of Schneider (2018), the visual system samples the visual scene, here a clock with rotating clock hand, during reoccurring intervals (“perceptual moments”, indicated by spaces between camera symbols). The end of these perceptual moments or “snapshots” (camera symbols) marks the registration of the stimulus position (the orientation of the clock hand and the flashed cue (red circle)). The stimuli are registered at the last known position. This is always the correct position for the clock hand, since it is moving at a fixed speed and constantly updated, but not for the transient red cue since the presentation dates back in time, and instantaneous updating cannot occur. Hence, a systematic lag of the red cue is perceived, its magnitude depending on the relative onset between cue and snapshot (camera symbol). In Panel A the red cue is presented very early in the perceptual moment. The temporal distance between the cue presentation and the end of the snapshot is large, leading to a long perceived flash lag. In Panel B the red cue is presented very shortly before the end of the perceptual moment. A short amount of time passes until the stimulus position is registered, and the perceived flash lag is brief.

Multiple explanations have been put forward to explain this phenomenon. We are going to briefly highlight the two most prominent ones, the differential latency theory (Whitney & Murakami, 1998) and the postdiction theory (Eagleman & Sejnowski, 2000). The differential latency hypothesis assumes that moving objects have a processing advantage over static objects and are therefore processed faster. The postdiction theory, on the other hand, proposes that position information is integrated for about 80 ms after the occurrence of the flash and is then used to compare the position of the two objects (for a more detailed comparison: Schneider, 2018).

A third hypothesis has been suggested, most prominently by Schneider (2018). He suggests that the FLE and other related illusions are a natural result of a discrete periodic sampling process. The theory states that visual information is collected over the time course of a so called “perceptual moment”. While information is collected continuously, the position of the object is registered only at the end of the perceptual moment and at its last known position. We can imagine a scenario where the static object is flashed right at the beginning of the perceptual moment (Figure 1A). The moving object would then move on for a specific period, until the end of the perceptual moment, at which the position of both objects is registered. In case A the perceived offset between moving and static object would be maximal. On the contrary, if the static object is flashed right at the end of the perceptual moment, the moving object would not move any further before the positions are registered, and the perceived offset would be minimal (Figure 1B). Based on the findings that the average perceived FLE is around 50 ms, with a standard deviation of 50 ms, the duration of a perceptual moment should be around 100 ms, which was verified by Schneider who investigated a large FLE dataset by Murakami (2001).

In this account, whether a long (Figure 1A) or short (Figure 1B) flash-lag illusion occurs on a given trial is mainly determined by the phase of the discrete sampling cycle at the moment of the flash onset. As this phase is generally unknown to the experimenter, the trial-to-trial variability in the illusion strength is often interpreted as noise. Some studies, however, have directly measured this phase with EEG, and verified that it influenced the flash-lag magnitude (Chakravarthi & Vanrullen, 2012). Here, our aim was to causally modulate the phase of the discrete sampling cycle by modulating the luminance of an annulus that surrounded a clock stimulus, and to prove that this phase had a causal influence on the flash-lag illusion. The FLE was randomly probed at 120 consecutive time points over the course of the entrainment (Figure 2). A frequency analysis of the average time course revealed a modulation of the perceived FLE duration at 10 Hz. We conclude that the visual stimulus entrained the discrete neural sampling mechanism, leading to a periodic modulation of the FLE.

**Figure 2.**
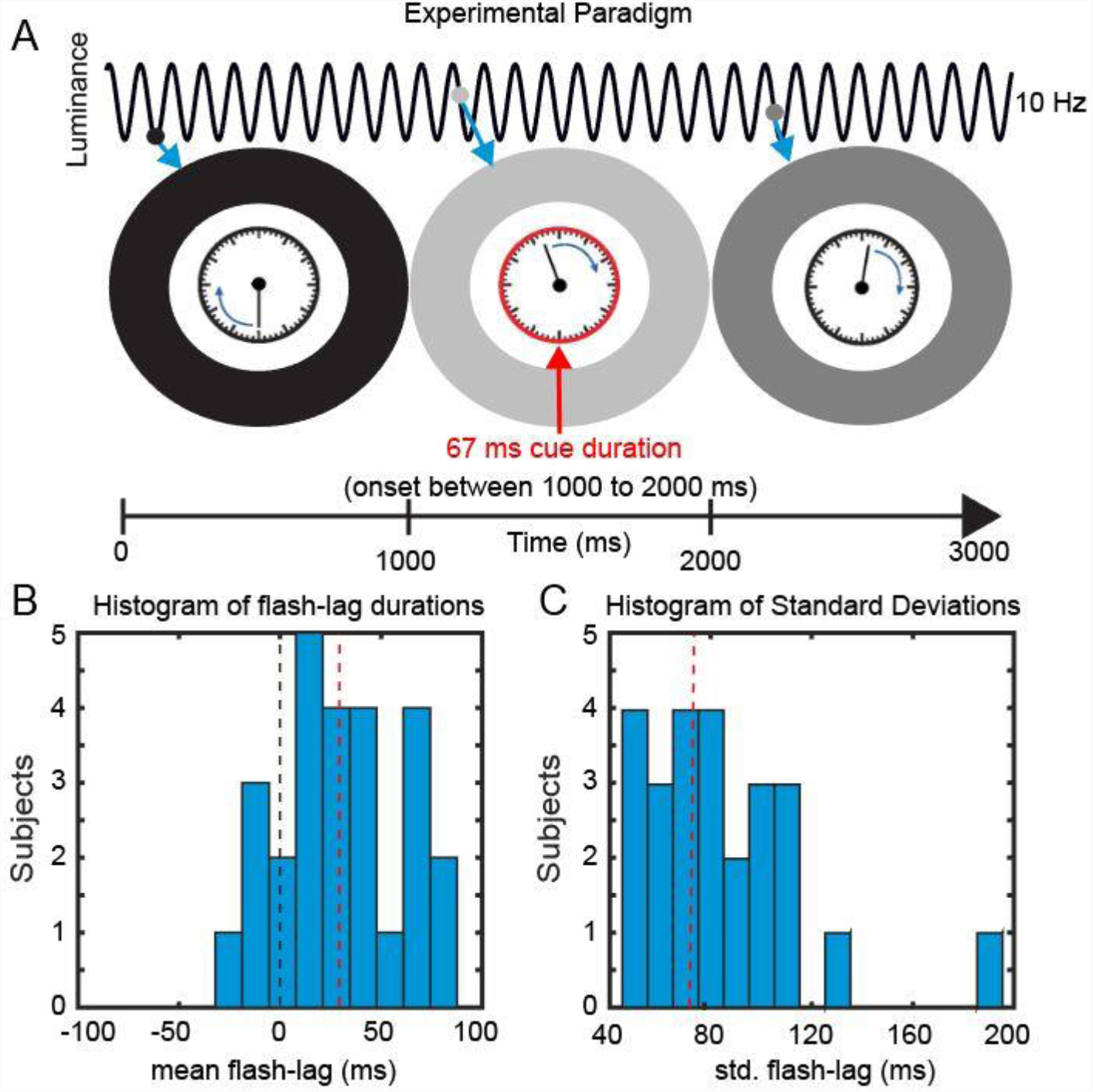
Experimental Paradigm. **A.** We presented a clock consisting of a static frame and a rotating clock hand for 3000 ms. The clock hand was rotating at 1 revolution per second and was surrounded by an entrainer annulus that periodically changed its luminance (from white to black) with a frequency of 10 Hz. At a random time point, within the central 1000 ms, a cue was presented in the form of the clock frame flashing red for 67 ms. Subsequently Participants reported the orientation of the clock hand at the onset of the cue. **B.** Behavioral Data. The mean perceived FLE duration was 32.4 ms (SEM across subjects: 6.48 ms). **C.** The mean std across trials was 75.4 ms (SEM across subjects: 6.42 ms).

## 2. Results

In the current study, we investigated the causal influence of a periodic entrainer on the perceived FLE duration. We presented participants with a clock stimulus containing a clock hand revolving at 1 Hz. The clock stimulus was surrounded by an entrainer annulus that changed its luminance from black to white periodically (following a sine function) at 10 Hz. At random SOAs we presented a cue by turning the frame of the clock red. Participants were then instructed to indicate the position of the clock hand at the onset of the Cue. The mean FLE duration (misperception in milliseconds) across observers was 32.4 ms (± 6.48 ms, SEM) (Figure 2B). Across trials the perceived FLE had an average standard deviation of 75.4 ms (±6.42 ms, SEM across subjects) (Figure 1C). Large standard deviations have been previously reported and have been shown to be unaffected by low level stimulus features (Chakravarthi & Vanrullen, 2012; Linares, Holcombe, & White, 2009). We were specifically interested in explaining this variability over trials in the context of a discrete sampling framework.

### 2.1 Visual entrainment modulates the perceived FLE

In order to verify an effect of the visual entrainment on the perceived FLE we calculated the average FLE time-series over individuals across the 120 SOAs. The original FLE time-series were down-sampled from 120 to 60 Hz, detrended by subtracting the 1 Hz Fourier component and normalized using a moving z-score, before averaging (window length 116 ms. See Methods section). Initial inspection of the time course (Figure 3A,B) indicates a strong oscillatory component, coupled with the background luminance modulation, in the 10 Hz range. To quantify this, we performed a frequency analysis on the preprocessed FL time-series. The resulting power spectrum revealed a dominant oscillation at 10 Hz (Figure 3C). To statistically test the significance of this peak we created 5000 surrogates by shuffling the 120 SOA-bin labels within subjects and recalculating the power spectrum. P-values were computed as the percentile of the mean power values within the bootstrapping distribution. This allowed us to test the null-hypothesis that the power spectrum of the average FLE time course does not show a peak at a specific frequency. All preprocessing steps were kept identical for the surrogates. The FLE time course oscillatory power at 10 Hz was significantly higher compared to the surrogate distribution (Figure 3C, p = 0.028, FDR corrected). We also analyzed the phase-consistency of the 10 Hz oscillation across subjects. The complex FFT coefficients at 10 Hz were extracted to calculate individual phase angles. We then compared these angles using Rayleigh’s test for non-uniformity testing the null hypothesis that the phase angles are randomly distributed. Phase angles were significantly clustered (Figure 3D, p < 0.05).

**Figure 3.**
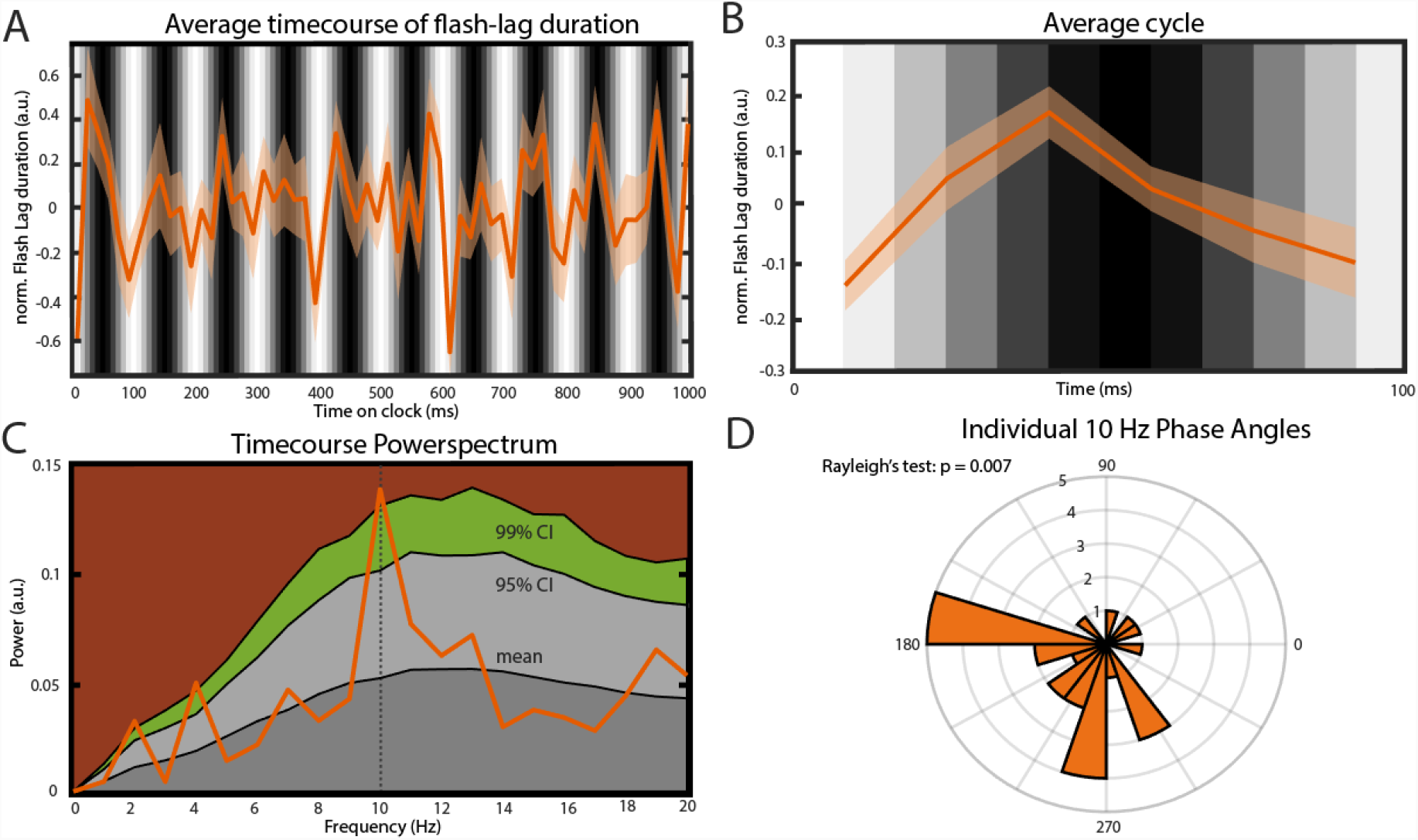
Main findings. **A.** Average time course of FLE (N = 25). Before averaging the individual time courses were down-sampled from 120 to 60 Hz, detrended by subtracting the 1 Hz Fourier component and normalized using a moving z-score (window length 100 ms. See Methods section.). The gray bars in the background indicate the luminance of the entrainer annulus at the moment of cue presentation. Transparent area shows the inter-subject SEM for the respective time point. **B.** Average cycle of the oscillation observed in A. Background Gray-scale bars indicate the luminance of the annulus at cue onset. Note that due to the down-sampling of the time course the 6 points of this average cycle always fall between two consecutive luminance bars. **C.** Power-spectrum of the average FLE time-course. The peak at 10 Hz was statistically compared to a surrogate distribution (5.000 surrogates) and was significant after correcting for multiple comparisons (p = 0.004, FDR-corrected: p = 0.028). Colored areas: Dark gray: Mean of the surrogate distribution; Light gray: 95% Confidence Interval; Green: 99% Confidence Interval; Brown: >99% Confidence Interval. **D.** Rose Plot of the 10 Hz Phases angles of individual FLE time-courses. The Rayleigh’s test of non-uniformity reveals a significant phase coherence between individual 10 Hz Phases (p = 0.007).

### 2.2 Control for annulus luminance

Figure 3 A and B show that bright luminance values tend to induce lower FLE values compared to dark luminance values. In accordance with our hypothesis, we interpret this effect as a result of the entrainment of the discrete sampling mechanism to the rhythmic luminance modulation. However, an alternative interpretation could be that the luminance of the background (even when it is not rhythmically modulated) has an effect on FLE. In order to control for the possible confound that the modulation in perceived FLE was caused merely by the luminance values of the annulus rather than neural entrainment caused by the dynamics of the stimulus, we conducted a control experiment (N = 25). The experimental parameters were kept identical to the dynamic entrainment condition with the exception of the annulus, which had a static luminance of 0% (black), 50% (gray) or 100% (white) throughout the trial. We statistically compared the perceived FLE in the three static luminance conditions using a one-way ANOVA (Figure 4). No significant difference in perceived FLE was observed between luminance conditions (F(2,1977) = 0.02, p = 0.98). We conclude that the luminance of the entrainer is not the main factor that explains the observed FLE modulation in the main experiment. Instead, it is likely that the rhythmic modulation of this luminance played a key role via rhythmic entrainment, in line with our hypothesis.

**Figure 4.**
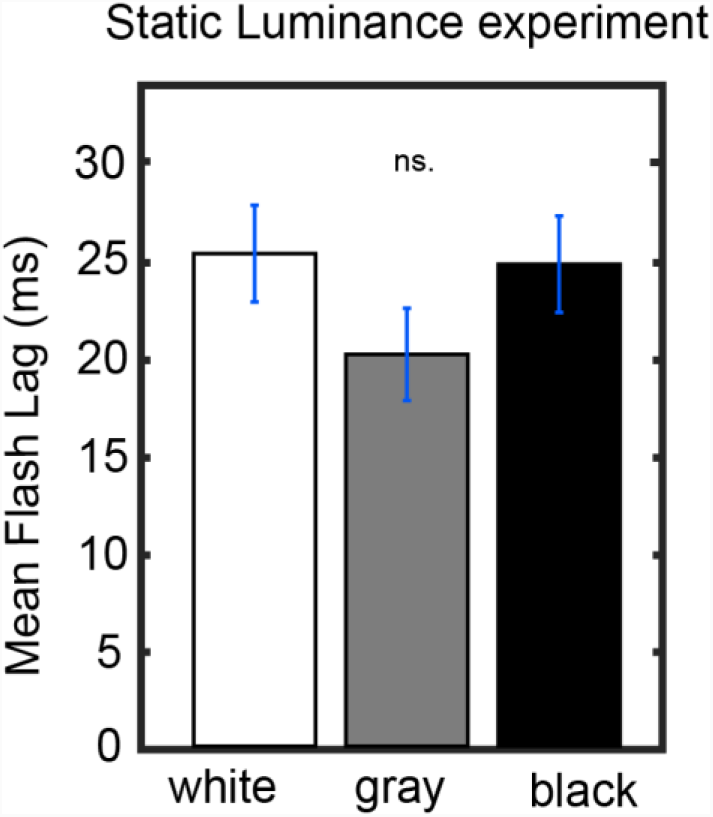
Control Experiment. Static luminance anuli were used to measure the effect of background luminance on the perceived FLE. The task paradigm was kept constant to that of the main experiment with the exception of the static annulus luminance. We did not find a significant effect of annulus luminance on the perceived FLE between any luminance conditions (one-way ANOVA, F(2,1977) = 0.02, p = 0.98).

## 3. Discussion

In this study we tested the causal influence of a visual entrainer at 10 Hz on the flash lag illusion. We found that the perceived FLE duration was periodically modulated at the entrainer frequency of 10 Hz and that these oscillations in the individual FLE time-series were strongly phase coherent between subjects. The oscillatory fluctuation in the perceived temporal offset between the moving and flashed object in the flash lag illusion is a direct demonstration of a rhythmic modulation of time perception. Our findings therefore provide strong causal evidence for the “hard” theory of discrete perception.

Our experiment was based on the discrete sampling hypothesis of perception (Busch et al., 2009; Haegens et al., 2011; Lőrincz et al., 2009; Samaha & Postle, 2015; Schneider, 2018; Valera et al., 1981; VanRullen, 2016; VanRullen & Koch, 2003; Vijayan & Kopell, 2012), which claims that the visual system periodically divides incoming visual information in discrete chunks. In the context of the flash lag illusion, visual information is assumed to be periodically collected over the course of a fixed time frame, sometimes called a “perceptual moment”, at the end of which the last known positions of the objects are registered (Schneider, 2018). If two objects are presented, one static and flashed and another moving continuously, a systematic offset between real and registered position is introduced, depending on the relative timing between presentation of the static object and the end of the perceptual moment (Figure 1).

The idea of a discrete sampling mechanism in vision affecting temporal perception has enjoyed a recent upswing in interest. Most recently it was shown that audio-visual entrainment at a participant’s individual alpha frequency influences performance in a visual integration/segregation task (Ronconi et al., 2018). Similarly, the phase of alpha oscillations can predict performance in a synchronous/asynchronous task (Milton & Pleydell-Pearce, 2016; Valera et al., 1981). Furthermore the frequency of individual alpha oscillations has been related to the participant’s two-flash fusion threshold, suggesting that visual systems with faster alpha rhythm sample the visual scene more frequently (Samaha & Postle, 2015). The above findings forward the existence of a certain integration window in the visual system within which information is merged. The fact that these windows can be dynamically modulated by rhythmic stimulation in psychophysical experiments as well as the correlational evidence provided by electrophysiological measurements hint at a neuro-oscillatory origin in the 8 to 12 Hz range, the alpha rhythm. Our findings fit well in this context, by providing causal evidence that a prominent temporal perceptual illusion can be dynamically modulated under the assumption of this discrete sampling mechanism.

What is the physiological explanation for our findings? The phase of alpha oscillations has been shown to be predictive of cortical excitability (Busch et al., 2009; Dugué, Marque, & VanRullen, 2011), of neuronal firing rates (Haegens et al., 2011; Lőrincz et al., 2009; Vijayan & Kopell, 2012) as well as of the amplitude of gamma oscillations (Osipova, Hermes, & Jensen, 2008; Voytek et al., 2010). As these neural signatures have been frequently implicated in neuronal processing it seems therefore logical that visual processing is concentrated on specific reoccurring intervals. The brain might use these naturally occurring periodicities, in the form of oscillations, to reduce the complexity of incoming information by compressing it into discrete packages. This compression might come at the cost of a reduced temporal resolution, leading to systematic errors in the perceived timing of stimuli as shown in this study (Schneider, 2018).

While our experimental paradigm was successful in eliciting the flash lag illusion, the mean perceived lag (32.4 ms) and the variation within participants (mean std: 75.4 ms) was different compared to some previous studies (Eagleman & Sejnowski, 2000; Kerzel, 2010; Murakami, 2001; Schneider, 2018). We attribute this discrepancy mainly to the paradigm that was applied in this experiment. Previous experiments have typically used only two visible stimuli to assess the FLE. In our paradigm multiple reference objects (i.e. minute markers) are on the screen that may aid the correct localization of the stimuli. An almost identical paradigm using the same clock stimulus was used by Chakravarthi and VanRullen (2012)and a highly similar mean FLE (27 ms) was observed, supporting this account. The mean std. across trials found here is somewhat closer to what has previously been reported. We think that the increase in variability might be a result of the highly dynamic entrainer annulus which might have had a distracting effect on attention. Without further experiments, however, we will not be able to explain these discrepancies fully.

Two important questions remain unanswered by our findings. First of all we only tested one specific frequency for the visual entrainment. While the alpha band has been previously related to the flash lag illusion (Chakravarthi & Vanrullen, 2012) and visual entrainment seems to be most effective at the brains endogenous alpha frequency (Herrmann, 2001), we cannot exclude the possibility that entrainment at a different frequency could lead to a similar modulation of the FLE. In order to address this question, this experiment should be repeated in the future, using different frequencies for entrainment. Second, it is unclear if a periodicity of the entraining stimulus is necessary. In our case it is unlikely but imaginable that transient annuli of white (or black) could be interpreted as single events by the brain, and lead to a subsequent dynamic modulation of the FLE. If this FLE modulation by each white (or black) transient was fast enough, this would become apparent as a 10 Hz oscillation in our FLE time-series. Unfortunately the two possibilities are close to impossible to disentangle, because single impulses, visual or applied with TMS, are often sufficient to elicit alpha oscillations as measured by EEG, and in turn would rhythmically influence the perceived FLE, which would also be consistent with our theory (Herring, Thut, Jensen, & Bergmann, 2015; VanRullen & Macdonald, 2012).

In this study, we successfully demonstrated that visual entrainment at 10 Hz leads to a periodic modulation of the FLE at an identical frequency. Furthermore this modulation was evident in most subjects, demonstrated by a strong inter-individual phase coherence of the individual FLE time courses. Our findings cannot be explained by static luminance states of the entrainer annulus, which we verified in a separate control experiment. In conclusion, we were able to provide causal evidence for the existence of a discrete sampling process in the visual system that gives rise to the flash lag illusion and can be dynamically modulated using visual entrainment. Our findings are in support of a “hard” version of discrete perception (whereby oscillations modulate not only sensory excitability, but also time perception), and highlight the involvement of the endogenous alpha rhythm of the brain.

## 4. Materials and Methods

### 4.1 Participants

26 participants (aged 18-30, 13 females) with normal or corrected to normal vision participated in the experiment. Informed consent forms were signed before the experiment. The experiment was carried out in accordance with the protocol approved by the Centre National de la Recherche Scientifique ethical committee and followed the Code of Ethics of the World Medical Association (Declaration of Helsinki).

### 4.2 Protocol

Stimuli were presented at a distance of 57 cm with a LCD display (1920 x 1080 resolution, 120 Hz refresh rate) using the Psychophysics Toolbox^i^ running in MATLAB (MathWorks). Stimuli consisted of a central fixation dot (diameter = 0.3°), a central clock stimulus (radius = 2°) with a black border (width = 0.3°), 60 evenly spaced clock markers (12 with length = 0.4°, 48 with length = 0.3°) and a rotating clock hand (length = 0.7°). The gap between the clock hand and the clock border was 1.3° and between clock hand and long clock marker 0.9°. The entrainer annulus and the clock hand were separated by 1.5°. The clock was surrounded by an entrainer annulus (outer radius = 11.5°, inner radius = 3.5°). Stimuli were presented on a gray background.

Trials started with the fixation point on the screen (Figure 2A). Participants initiated the trial via button press. Directly after the button press the entrainer annulus as well as the clock stimulus appeared on the screen with the clock hand rotating with 1 rotation/s. The luminance of the entrainer annulus was modulated sinusoidally at 10 Hz with the luminance ranging from 0 to 255 (0.68 to 100.8 cd/mm^2^). At a random time point between 1000 ms and 2000 ms (SOA window) a cue was presented (the frame of the clock stimulus turned red) for 66 ms (8 frames). Afterwards the clock hand continued rotating until the 3000 ms mark was reached. After a delay of 1000 ms an identical clock with a static hand was presented. The participant could rotate the clock hand with the arrow keys to indicate the perceived location of the clock hand at the time point of Cue onset. The Cue onset was randomly chosen from a discrete uniform distribution of the 120 SOAs between 1000 ms and 2000 ms. Participants each performed 480 trials resulting in 4 responses per SOA. Participants were instructed to maintain central fixation during the 3 second window when the clock was on the screen. The starting position of the clock hand was randomized between subjects but was kept constant for a single subject.

### 4.3 Control experiment

We conducted a control experiment to verify that the observed modulation was due to neural entrainment and not simply due to the luminance of the annulus. A new set of participants (N=25) conducted a similar experiment where the luminance of the annulus was kept constant throughout the trial. Instead of testing all 7 luminance values we tested the most extremes ones (black and white) as well as 50% luminance at which the annulus is indistinguishable from the background. In the original experiment 40 trials were collected for each of the two extreme luminance values and 80 trials for the 50% luminance per participant. In the control experiment we collected 40 trials for each of the three luminance values resulting in 120 trials per subject. To compare the perceived FLE between the luminance values in the control experiment a one-way ANOVA was conducted.

### 4.4 Data Analysis

Attributing one SOA (frame 1 to frame 120) to every position of the clock hand we can define the FLE duration as the temporal distance (in frames) between the actual onset of the Cue (position of the clock hand when clock border turns red) and the orientation of the clock hand that the participant indicated. Note that we define SOA as the temporal distance to the onset of the central 1 s window, within which the Cue could appear. We restricted our analysis on responses that were within 3.5 times the standard deviation of each individual participant (mean: 290.2 ms, SEM: +-22.3 ms) before or after the actual Cue onset. Single trial responses were averaged over the respective SOA (4 per SOA, 120 SOAs). The resulting time course represents the average FLE as a function of time for one individual. The individual time course was then down-sampled from 120 to 60 Hz. To eliminate effects of the position of the clock hand on the flash lag illusion, we fitted a 1 Hz sinus function and subtracted it from the individual time course. The frequency of the fit was determined by assuming that any effects stemming from the clock hand position should influence the flash lag illusion at a frequency identical to the revolution frequency 1 Hz. We then normalized the data by applying a moving z-score window of length 116 ms. A 116 ms window (7 SOAs) of the original data was z-scored and the central value was saved in a separate array. The window was then shifted by 16 ms (1 SOA) and the process was repeated resulting in one normalized time course per subject. We validated in a separate re-analysis that the length of the window did not affect our findings. Window lengths of 60 ms up to 208 ms lead to comparable modulations at 10 Hz. Individual time courses were then averaged and analyzed in the frequency domain using FFT. 60 SOAs at 60 Hz allowed for a Nyquist frequency of 30 Hz. Only frequencies from 1 Hz to 2/3 of the Nyquist frequency (1 to 20 Hz, 20 values) were considered. The complex FFT coefficients were squared to obtain oscillatory power at each frequency. To statistically test if the power at 10 Hz is significant we calculated 5000 surrogates by shuffling the SOA-labels between trials, and repeating all analysis steps for each surrogate as explained above (60Hz down-sampling, 1Hz detrending, normalization). The original power-spectrum was then compared to the surrogate distribution and p-values were corrected for multiple comparisons using the False Discovery Rate. Individual Phase angles were extracted from the 10 Hz component of the FFT of the down-sampled and normalized time-courses. Rayleigh’s test for non-uniformity was used to statistically test if individual phases were significantly coherent.

## Author Contribution Statement

S.C. designed and conducted the experiment, analyzed the results and wrote the manuscript. R.V. edited the manuscript. All authors provide their approval for submission

## References

Ai, L., & Ro, T. (2013). The phase of prestimulus alpha oscillations affects tactile perception. Journal of Neurophysiology, 111(6), 1300–1307. https://doi.org/10.1152/jn.00125.2013

Başar, E., Başar-Eroglu, C., Karakaş, S., & Schürmann, M. (2001). Gamma, alpha, delta, and theta oscillations govern cognitive processes. International Journal of Psychophysiology: Official Journal of the International Organization of Psychophysiology, 39(2–3), 241–248.

Busch, N. A., Dubois, J., & VanRullen, R. (2009). The phase of ongoing EEG oscillations predicts visual perception. The Journal of Neuroscience: The Official Journal of the Society for Neuroscience, 29(24), 7869–7876. https://doi.org/10.1523/JNEUROSCI.0113-09.2009

Callaway, E., & Yeager, C. L. (1960). Relationship between reaction time and electroencephalographic alpha phase. Science (New York, N.Y.), 132(3441), 1765–1766.

Chakravarthi, R., & Vanrullen, R. (2012). Conscious updating is a rhythmic process. Proceedings of the National Academy of Sciences of the United States of America, 109(26), 10599–10604. https://doi.org/10.1073/pnas.1121622109

Dugué, L., Marque, P., & VanRullen, R. (2011). The phase of ongoing oscillations mediates the causal relation between brain excitation and visual perception. The Journal of Neuroscience: The Official Journal of the Society for Neuroscience, 31(33), 11889–11893. https://doi.org/10.1523/JNEUROSCI.1161-11.2011

Dugué, L., & VanRullen, R. (2017). Transcranial Magnetic Stimulation Reveals Intrinsic Perceptual and Attentional Rhythms. Frontiers in Neuroscience, 11. https://doi.org/10.3389/fnins.2017.00154

Eagleman, D. M., & Sejnowski, T. J. (2000). Motion Integration and Postdiction in Visual Awareness. Science, 287(5460), 2036–2038. https://doi.org/10.1126/science.287.5460.2036

Haegens, S., Nácher, V., Luna, R., Romo, R., & Jensen, O. (2011). α-Oscillations in the monkey sensorimotor network influence discrimination performance by rhythmical inhibition of neuronal spiking. Proceedings of the National Academy of Sciences of the United States of America, 108(48), 19377–19382. https://doi.org/10.1073/pnas.1117190108

Herring, J. D., Thut, G., Jensen, O., & Bergmann, T. O. (2015). Attention Modulates TMS-Locked Alpha Oscillations in the Visual Cortex. The Journal of Neuroscience: The Official Journal of the Society for Neuroscience, 35(43), 14435–14447. https://doi.org/10.1523/JNEUROSCI.1833-15.2015

Herrmann, C. S. (2001). Human EEG responses to 1–100 Hz flicker: resonance phenomena in visual cortex and their potential correlation to cognitive phenomena. Experimental Brain Research, 137(3), 346–353. https://doi.org/10.1007/s002210100682

Keitel, C., Quigley, C., & Ruhnau, P. (2014). Stimulus-Driven Brain Oscillations in the Alpha Range: Entrainment of Intrinsic Rhythms or Frequency-Following Response? Journal of Neuroscience, 34(31), 10137–10140. https://doi.org/10.1523/JNEUROSCI.1904-14.2014

Kerzel, D. (2010). The Fröhlich effect: past and present. In R. Nijhawan & B. Khurana (Eds.), Space and Time in Perception and Action (pp. 321–337). Cambridge: Cambridge University Press. https://doi.org/10.1017/CBO9780511750540.019

Linares, D., Holcombe, A. O., & White, A. L. (2009). Where is the moving object now? Judgments of instantaneous position show poor temporal precision (SD = 70 ms). Journal of Vision, 9(13), 9–9. https://doi.org/10.1167/9.13.9

Lőrincz, M. L., Kékesi, K. A., Juhász, G., Crunelli, V., & Hughes, S. W. (2009). Temporal Framing of Thalamic Relay-Mode Firing by Phasic Inhibition during the Alpha Rhythm. Neuron, 63(5), 683–696. https://doi.org/10.1016/j.neuron.2009.08.012

Mathewson, K. E., Fabiani, M., Gratton, G., Beck, D. M., & Lleras, A. (2010). Rescuing stimuli from invisibility: Inducing a momentary release from visual masking with pre-target entrainment. Cognition, 115(1), 186–191. https://doi.org/10.1016/j.cognition.2009.11.010

Mathewson, K. E., Gratton, G., Fabiani, M., Beck, D. M., & Ro, T. (2009). To See or Not to See: Prestimulus α Phase Predicts Visual Awareness. Journal of Neuroscience, 29(9), 2725–2732. https://doi.org/10.1523/JNEUROSCI.3963-08.2009

Mathewson, K. E., Lleras, A., Beck, D. M., Fabiani, M., Ro, T., & Gratton, G. (2011). Pulsed Out of Awareness: EEG Alpha Oscillations Represent a Pulsed-Inhibition of Ongoing Cortical Processing. Frontiers in Psychology, 2. https://doi.org/10.3389/fpsyg.2011.00099

Milton, A., & Pleydell-Pearce, C. W. (2016). The phase of pre-stimulus alpha oscillations influences the visual perception of stimulus timing. NeuroImage, 133, 53–61. https://doi.org/10.1016/j.neuroimage.2016.02.065

Murakami, I. (2001). A flash-lag effect in random motion. Vision Research, 41(24), 3101–3119. https://doi.org/10.1016/S0042-6989(01)00193-6

Osipova, D., Hermes, D., & Jensen, O. (2008). Gamma power is phase-locked to posterior alpha activity. PloS One, 3 (12), e3990. https://doi.org/10.1371/journal.pone.0003990

Romei, V., Gross, J., & Thut, G. (2010). On the Role of Prestimulus Alpha Rhythms over Occipito-Parietal Areas in Visual Input Regulation: Correlation or Causation? Journal of Neuroscience, 30(25), 8692–8697. https://doi.org/10.1523/JNEUROSCI.0160-10.2010

Ronconi, L., Busch, N. A., & Melcher, D. (2018). Alpha-band sensory entrainment alters the duration of temporal windows in visual perception. Scientific Reports, 8(1), 11810. https://doi.org/10.1038/s41598-018-29671-5

Samaha, J., & Postle, B. R. (2015). The Speed of Alpha-Band Oscillations Predicts the Temporal Resolution of Visual Perception. Current Biology, 25(22), 2985–2990. https://doi.org/10.1016/j.cub.2015.10.007

Schneider, K. A. (2018). The Flash-Lag, Fröhlich and Related Motion Illusions Are Natural Consequences of Discrete Sampling in the Visual System. Frontiers in Psychology, 9. https://doi.org/10.3389/fpsyg.2018.01227

Spaak, E., Lange, F. P. de, & Jensen, O. (2014). Local Entrainment of Alpha Oscillations by Visual Stimuli Causes Cyclic Modulation of Perception. Journal of Neuroscience, 34(10), 3536–3544. https://doi.org/10.1523/JNEUROSCI.4385-13.2014

Thut, G., Veniero, D., Romei, V., Miniussi, C., Schyns, P., & Gross, J. (2011). Rhythmic TMS Causes Local Entrainment of Natural Oscillatory Signatures. Current Biology, 21(14), 1176–1185. https://doi.org/10.1016/j.cub.2011.05.049

Valera, F. J., Toro, A., Roy John, E., & Schwartz, E. L. (1981). Perceptual framing and cortical alpha rhythm. Neuropsychologia, 19(5), 675–686. https://doi.org/10.1016/0028-3932(81)90005-1

VanRullen, R. (2016). Perceptual Cycles. Trends in Cognitive Sciences, 20(10), 723–735. https://doi.org/10.1016/j.tics.2016.07.006

VanRullen, R., & Koch, C. (2003). Is perception discrete or continuous? Trends in Cognitive Sciences, 7(5), 207–213.

VanRullen, R., & Macdonald, J. S. P. (2012). Perceptual echoes at 10 Hz in the human brain. Current Biology: CB, 22(11), 995–999. https://doi.org/10.1016/j.cub.2012.03.050

Vijayan, S., & Kopell, N. J. (2012). Thalamic model of awake alpha oscillations and implications for stimulus processing. Proceedings of the National Academy of Sciences, 109(45), 18553–18558. https://doi.org/10.1073/pnas.1215385109

Voytek, B., Canolty, R. T., Shestyuk, A., Crone, N. E., Parvizi, J., & Knight, R. T. (2010). Shifts in Gamma Phase–Amplitude Coupling Frequency from Theta to Alpha Over Posterior Cortex During Visual Tasks. Frontiers in Human Neuroscience, 4. https://doi.org/10.3389/fnhum.2010.00191

Whitney, D., & Murakami, I. (1998). Latency difference, not spatial extrapolation. Nature Neuroscience, 1(8), 656–657. https://doi.org/10.1038/3659

